# Genomic sequence analysis of the first mpox virus detected in Kenya

**DOI:** 10.1101/2024.08.20.608891

**Authors:** Solomon K. Langat, Konongoi Limbaso, Samoel Khamadi, Albert Nyunja, Genay Pilarowski, Emmanuel Okunga, Victor Ofula, Paul Oluniyi, Hellen Koka, Edith Koskei, Samuel Owaka, Betty Chelangat, Francis Mulwa, James Mutisya, Edith Chepkorir, Joel Lutomiah, Daniel Langat, Patrick Amoth, Elijah Songok

**Author notes:** These authors contributed equally to this work (Solomon K. Langat, Konongoi Limbaso).

## Abstract

Mpox is a zoonotic disease caused by the Monkeypox virus (MPXV) in the family: Poxviridae, genus: Orthopoxvirus. Historically, the disease was restricted mostly to Africa with cases being reported in Central Africa (mostly caused by clade I) and in West Africa (caused by clade II). However, there has been a recent shift in the virus range with outbreaks being reported in Europe and America, and in countries where the virus was initially not endemic. This multi-country outbreak was driven mostly by clade IIb lineage of MPXV. Since December 2023, there has been an ongoing mpox outbreak in the Democratic Republic of Congo (DRC), driven by a new clade I lineage of the virus, designated clade Ib. The DRC outbreak has persisted, with an increase in cases being reported over the past few months. Spillover of these outbreak-related cases to the neighbouring countries have also been reported in multiple countries including Uganda and Rwanda. Here, we report the rapid application of unbiased metagenomic next generation sequencing (mNGS) to reconstruct the genome sequence of the first reported case of MPXV in Kenya. Our findings show that the Kenyan case clusters together with clade Ib MPXV strains, associated with the sustained outbreak in the DRC. Clade Ib lineage has been associated with continuing geographical expansion of the virus to previously unaffected areas, high incidence of the disease as well as high case fatality (CFR 4.9-6.7%). Similar to other clade Ib strains, the Kenyan strain carries predominant APOBEC3-type mutations which is characteristic feature of human-to-human transmission, highlighting the need for surveillance to curtail any potential expansion of this MPXV strain. The lack of information on genomes associated with cases reported in different East African countries, is a gap that urgently needs to be addressed to aid in the monitoring of this MPXV strain. This case investigation, therefore, underscores the need for sequencing efforts to be enhanced across the continent to help improve our understanding of the geographical range and diversity of the MPXV strains, especially those belonging to clade I which is currently under-represented.

## Introduction

Mpox is a zoonotic disease caused by monkeypox virus (MPXV), belonging to the Orthopoxvirus genus in the family *Poxviridae* (1). It was first discovered in 1958 among laboratory research primates, but it was not until 1970 when the first case of MPXV was detected in a 9-month-old child in the Democratic Republic of Congo (DRC) (2,3). This was followed by decades of mpox case detections that were restricted mostly to selected African countries, particularly the DRC. MPXV is large, brick-shaped and enveloped double-stranded linear DNA (dsDNA) genome that is approximately 190kb (10). The virus particle is extremely complex containing more than 100 viral proteins. The viral genome has roughly 200 open reading frames which are transcribed at different times in the early, intermediate, and late stages of infection (10). Phylogenetic analyses have placed the virus into two major clades, clade I (Central African) and Clade II (West African) (11). Clade I is common in Central Africa and has been associated with severe clinical symptoms and higher mortality rates (CFR 4-11%) (13,20). On the other hand, clade II was initially confined to West Africa and causes less severe symptoms and mortality (CFR <4%) (13,20).

Historically, Clade I MPXV has been the most common lineage associated with majority of the reported mpox cases (up to 95% of the cases). However, since May 2022 there has been a shift in the virus range with clade II lineage of the MPXV causing a global epidemic that led to case detections in Europe and America, and in countries where the virus was initially not endemic (4). Several African countries have also reported cases of MPXV since 2022. Kenya, however, did not record any case during this multi-country outbreak. Epidemiological studies revealed that most of the cases reported globally were driven by MPXV clade IIb and were transmitted through sexual contact (4,15,16). Since December 2023, there has been an ongoing mpox outbreak in the DRC, driven by a new clade I lineage of the virus designated clade Ib (5,6). The DRC outbreak has persisted, with increase in cases being reported over the past few months (6,7,8). Spillover of these outbreak-related cases to the neighbouring countries have also been reported in multiple countries including Uganda and Rwanda. Following this sustained outbreak, and the expanding geographical range of the virus including to the neighbouring countries, World Health Organization and the Africa CDC recently declared the outbreak as a public health emergency of continental and international concern (17,18). Genomic analyses of this new clade of MPXV, clade Ib, revealed a mutational pattern which suggests an evolution of the virus driven by an apolipoprotein B messenger RNA editing enzyme, catalytic subunit 3 (APOBEC3) cytosine deamination (8). This has previously been shown to be a characteristic feature of human-to-human transmission of MPXV (14). The new clade I lineage (clade Ib) has been associated with continuing geographical expansion to previously unaffected areas, high incidence of the disease as well as high case fatality (CFR 4.9-6.7%) (8,9). Approximately 88% of the mortalities reported in the ongoing DRC outbreak have been associated with children younger than 15 years of age, from a case load of about 70% of the total reported cases (7).

Sequencing efforts have been key to understanding the genomic characteristics and epidemiology of MPXV, allowing for implementation of tailored control strategies. These genomic efforts allow for the identification of the causative virus strain, understanding its evolutionary trajectory, as well as genetic and phenotypic characteristics which are critical in guiding diagnostics as well as vaccine development. Here, we report the rapid application of unbiased metagenomic next generation sequencing (mNGS) to reconstruct the genome sequence of a reported case of mpox in Kenya.

## Methods

### The case

The reported case was from a 42-year-old male, a long-distance truck driver who had travelled from Kampala, Uganda to Mombasa, Kenya and was enroute to Rwanda at the time of detection. The date of onset of symptoms was captured as 3^rd^ July 2024. The case was detected and isolated by the Kenyan Ministry of Health officials on 22^nd^ July 2024 after presenting with classic symptoms of mpox infection that included septic spots on the face, neck, chest, back, forearms, private parts and feet. The sample consisting of lesion material was obtained by the Ministry of Health officials on 25^th^ July 2024 in the form of skin lesion material and referred to KEMRI for testing. The protocol under which the sample was obtained and processed for quantitative real-time PCR (qPCR) and sequencing is approved by the Kenya Medical Research Institute’s Scientific Ethics and Review Unit (SERU) under study number; SSC3035.

### Molecular characterisation

#### Extraction

DNA extraction was carried out on the clinical sample (lesions and vesicle swabs) using Biocomma DNA/RNA virus purification kit. MPXV was detected on the extracted DNA using the Liferiver Real Time PCR Kit, following the manufacturer’s instructions. The DNA was subsequently subjected to unbiased metagenomic next-generation sequencing (mNGS). Libraries were first prepared using NEBNext Ultra II DNA library prep kit for Illumina (NEB, UK), and the prepared libraries were sequenced on an Illumina iSeq 100 to generate a 2×146bp paired-end reads.

#### Sequence Analysis

Raw sequence reads were submitted to CZ ID (https://czid.org/), which is an integrated pipeline that allows de-hosting, quality control and adapter trimming as well as assembly of the cleaned paired reads. *De novo*-generated contigs were used to carry out preliminary analysis of the MPXV strain, including determining the lineage of the virus using Nextclade (19). Following the preliminary placement of the MPXV strain to clade I, we utilized CZ ID viral consensus genome pipeline in order to generate a consensus genome from the sequence reads using an earlier clade I lineage MPXV genome (accession NC_003310) as reference. To further investigate the sequenced isolate, in the context of other global outbreak-associated strains, we performed phylogenetic analysis using sequences downloaded from Genbank (supplementary material). We selected all high-quality sequences (minimum length >160k) from Africa, South America, Asian continent and Oceania. Due to their high representation, sequences from Europe and North America were randomly subsampled using the NCBI Virus randomization option, such that a maximum of 20 sequences from each country were selected for inclusion. Only sequences with complete metadata information, including geographical location and collection date, were selected for phylogenetic analysis. The combined set of sequences were analysed using squirrel bioinformatics pipeline (https://github.com/aineniamh/squirrel). The pipeline performs multiple sequence alignment, masking of low-complexity regions that have been characterized for MPXV, maximum-likelihood phylogenetic analysis and ancestral state reconstruction (ASR) to characterize APOBEC3-like and non-APOBEC3 mutations. The phylogenetic tree generated was visualized in Figtree v1.4.4 (https://github.com/rambaut/figtree). The sequencing data generated can be found in NCBI BioProject PRJNA1147890 and the genome sequence reported in the study has been submitted to Genbank under accession number PQ178862.

## Results and Discussion

DNA extraction was performed on the sample received, and it was confirmed as positive for MPXV by PCR with a cycling threshold (Ct) of ∼20. We subsequently subjected the residual DNA to unbiased mNGS. A total of 6.3 million reads were generated from the run. Approximately 99% of the total reads recovered were dropped after removal of the host reads, low quality reads and duplicates. The remaining sequence reads (∼74,500) were used for viral genome assembly. We assembled a near-complete genome with a total length of approximately 189kb, which is equivalent to approximately 94% of the entire MPXV genome. The average depth of coverage achieved was 20.3x. Comparison to other MPXV genomes available in Genbank showed that the sequenced strain had a 99.5% nucleotide similarity to MPXV sequences from the DRC. Phylogenetic analysis placed the MPXV associated with the Kenyan case in clade I, clustering together with genomes associated with the 2024 DRC outbreak (Figure1). This outbreak clade forms a unique cluster separate from the previous clade I strains, and it has been proposed as a new subclade, clade Ib, with the earlier strains being proposed as clade Ia (Figure1) (5). Ancestral state reconstruction identified 7 unique single nucleotide polymorphisms (SNPS) carried by the Kenyan strain. Five of these were APOBEC3-like mutations, consistent with earlier finding that clade Ib had predominance of APOBEC3-type mutations which indicates its adaptation to circulation among humans (8). The new variant, clade Ib, has been associated with sustained human-to-human transmission in the South Kivu province of the DRC (8). The sustained outbreak linked to MPXV clade Ib variant, as well as the associated high incidence and case fatality raises concerns of its transmissibility as well as the disease severity. Further, there has been continuing geographic expansion of this outbreak to previously unaffected areas (8). Recent reports of detection of MPXV in several neighbouring countries such as Rwanda and Uganda, and now the confirmation of this variant in Kenya points to continuing risk of expansion to the neighbouring countries and highlights the need for vigilance to curtail further geographic expansion of this MPXV strain.

**Figure1:**
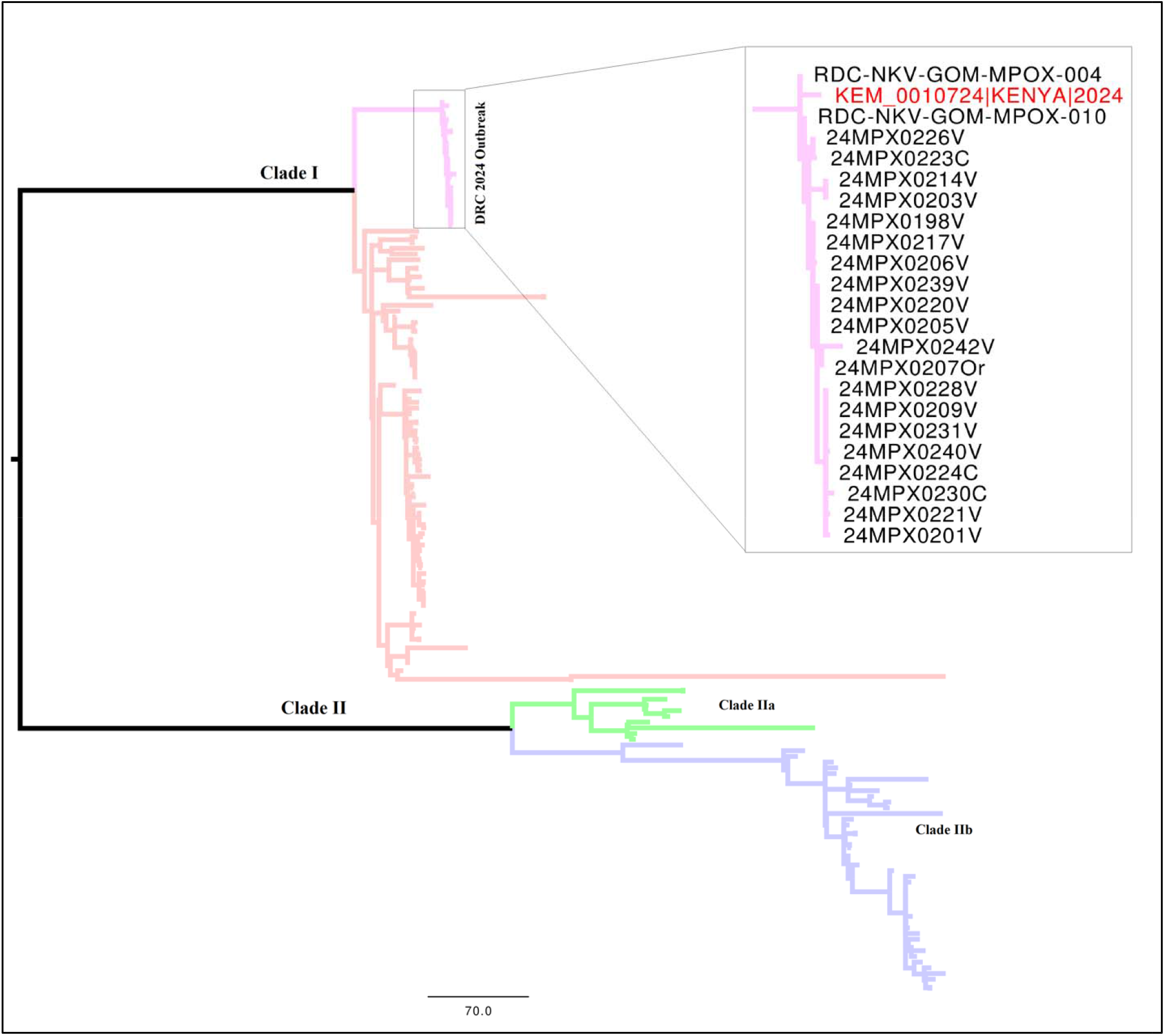
Maximum Likelihood phylogeny based on complete genomes belonging to the two major clades of MPXV. The DRC outbreak clade has-been expanded with the sequenced strain from Kenya indicated with a red tip label.

## Conclusion

We performed unbiased DNA mNGS on a case of MPXV in Kenya. The investigation recovered a near-complete genome of the virus, and our analysis shows that it belongs to clade Ib. The current strain clusters closely with MPXV strains recently associated with sustained human-to-human transmission in the DRC. Similar to other clade Ib strains, the Kenyan strain carries predominant APOBEC3-type mutations. Our findings highlight the need for continued surveillance to curtail any expansion of this MPXV strain in Kenya and the region. The lack of information on genomes from cases reported in different East African countries, is a gap that urgently needs to be addressed to aid in the identification of risk factors and inform mitigation strategies. There is, therefore, need for sequencing efforts to be enhanced across the continent to help improve our understanding of the geographical range and diversity of the MPXV strains, especially those belonging to clade I which is currently under-represented in public databases.

## Acknowledgments

We wish to thank all the hospital personnel and the Disease Surveillance and Response Unit for identifying the suspect case, facilitating the collection and shipment of samples for testing and reporting to guide the public health response measures.

This work was made possible through the support of KEMRI, Bill and Melinda Gates Foundation and Chan Zuckerberg Initiative (CZI) grant INV-050635. The findings and conclusions in this report are those of the authors and do not necessarily represent the official position of the funders.

