## Supplementary material for "Genomic sequence analysis of the first mpox virus detected in Kenya"

List_of_Genbank_mpox_sequences

| strain | accession | date_submitted | date_of_collection | country | clade_membership | author |
|---|---|---|---|---|---|---|
| 2091 | OL504741 | 2022-07-06 | 2019-12-XX | United Kingdom | IIb | Atkinson,B.,Pottage,T.,Ngabo,D.,Crook,A.,Pitman,J.,Summers,S.,Pullan,S.,Lewandowski,K.,Furneaux,J.,Davies,K.,Brooks,T. |
| 2102 | OL504743 | 2022-07-06 | 2019-12-XX | United Kingdom | IIb | Atkinson,B.,Pottage,T.,Ngabo,D.,Crook,A.,Pitman,J.,Summers,S.,Pullan,S.,Lewandowski,K.,Furneaux,J.,Davies,K.,Brooks,T. |
| KC257459 | KC257459 | 2013-02-20 | 2005-XX-XX | Sudan | I | Nakazawa,Y.,Emerson,G.L.,Carroll,D.S.,Zhao,H.,Li,Y.,Reynolds,M.G.,Karem,K.L.,Olson,V.A.,Lash,R.R.,Davidson,W.B.,Smith,S.K.,Levine,R.S.,Regnery,R.L.,Sammons,S.A.,Frace,M.A.,Mutasim,E.M.,Karsani,M.E.,Muntasir,M.O., |
| KC257460 | KC257460 | 2013-02-20 | 1985-XX-XX | Democratic Republic of the Congo | I | Nakazawa,Y.,Emerson,G.L.,Carroll,D.S.,Zhao,H.,Li,Y.,Reynolds,M.G.,Karem,K.L.,Olson,V.A.,Lash,R.R.,Davidson,W.B.,Smith,S.K.,Levine,R.S.,Regnery,R.L.,Sammons,S.A.,Frace,M.A.,Mutasim,E.M.,Karsani,M.E.,Muntasir,M.O., |
| Zaire 1979-005 | HM172544 | 2010-07-25 | 1979-XX-XX | Democratic Republic of the Congo | I | Farlow,J.,Ichou,M.A.,Huggins,J.,Ibrahim,S.,Ait Ichou,M.,Ibrahims,S. |
| Zaire_1979-005 | DQ011155 | 2005-09-27 | 1978-XX-XX | Democratic Republic of the Congo | I | Likos,A.M.,Sammons,S.A.,Olson,V.A.,Frace,A.M.,Li,Y.,Olsen-Rasmussen,M.,Davidson,W.,Galloway,R.,Khristova,M.L.,Reynolds,M.G.,Zhao,H.,Carroll,D.S.,Curns,A.,Formenty,P.,Esposito,J.J.,Regnery,R.L.,Damon,I.K.,Olsen-Ras |
| V79-I-005 | HQ857562 | 2011-02-15 | 1979-XX-XX | Democratic Republic of the Congo | I | Estep,R.D.,Messaoudi,I.,O'Connor,M.A.,Li,H.,Sprague,J.,Barron,A.,Engelmann,F.,Yen,B.,Powers,M.F.,Jones,J.M.,Robinson,B.A.,Orzechowska,B.U.,Manoharan,M.,Legasse,A.,Planer,S.,Wilk,J.,Axthelm,M.K.,Wong,S.W. |
| MN702446 | MN702446 | 2020-08-29 | 2018-04-14 | Central African Republic | I | Selekon,B.,Labouba,I.L.,Gonofio,E.C.,Sem Ouilibona,R.,Simo Tchetgna,H.,Besombes,C.,Feher,M.,Fontanet,A.,Kazanji,M.,Manugerra,J.-C.,Gessain,A.,Nakoune,E.,Berthet,N.,Nkili Meyong,A. |
| MN702449 | MN702449 | 2020-08-29 | 2016-01-02 | Central African Republic | I | Selekon,B.,Labouba,I.L.,Gonofio,E.C.,Sem Ouilibona,R.,Simo Tchetgna,H.,Besombes,C.,Feher,M.,Fontanet,A.,Kazanji,M.,Manugerra,J.-C.,Gessain,A.,Nakoune,E.,Berthet,N.,Nkili Meyong,A. |
| MT724772 | MT724772 | 2021-03-28 | 2014-01-01 | Democratic Republic of the Congo | I | Marien,J.,Laudisoit,A.,Patrono,L.V.,Calvignac-Spencer,S.,Leendertz,F.,Leirs,H.,Verheyen,E. |
| MT724770 | MT724770 | 2021-03-28 | 2014-01-01 | Democratic Republic of the Congo | I | Marien,J.,Laudisoit,A.,Patrono,L.V.,Calvignac-Spencer,S.,Leendertz,F.,Leirs,H.,Verheyen,E. |
| MN702445 | MN702445 | 2020-08-29 | 2017-02-06 | Central African Republic | I | Selekon,B.,Labouba,I.L.,Gonofio,E.C.,Sem Ouilibona,R.,Simo Tchetgna,H.,Besombes,C.,Feher,M.,Fontanet,A.,Kazanji,M.,Manugerra,J.-C.,Gessain,A.,Nakoune,E.,Berthet,N.,Nkili Meyong,A. |
| MN702444 | MN702444 | 2020-08-29 | 2017-02-06 | Central African Republic | I | Selekon,B.,Labouba,I.L.,Gonofio,E.C.,Sem Ouilibona,R.,Simo Tchetgna,H.,Besombes,C.,Feher,M.,Fontanet,A.,Kazanji,M.,Manugerra,J.-C.,Gessain,A.,Nakoune,E.,Berthet,N.,Nkili Meyong,A. |
| KJ642613 | KJ642613 | 2015-05-11 | 1970-XX-XX | Democratic Republic of the Congo | I | Nakazawa,Y.,Mauldin,M.R.,Emerson,G.L.,Reynolds,M.G.,Lash,R.R.,Gao,J.,Zhao,H.,Li,Y.,Muyembe,J.J.,Kingebeni,P.M.,Wemakoy,O.,Malekani,J.,Karem,K.L.,Damon,I.K.,Carroll,D.S.,Lil,Y.,Weil,R.M.,Muyembe,J.-J.,Kingbeni,P.M |
| 24MPX0008V | PP601188 | 2024-05-02 | 2023-12-16 | Democratic Republic of the Congo | I | Vakaniaki,E.H.,Kaciat,C.,Kinganda - Lusamaki,E.,O'Toole,A.,Wawina -Bokalanga,T.,Mukadi - Bamuleka,D.,Amuri,A.A.,Parker,E.,Muswamba-Kayembe,P.-C.,Makangara - Cigolo,J.-C.,Mulopo - Mukanya,N.,Pukuta - Simbu,E.,M |
| MN702451 | MN702451 | 2020-08-29 | 2017-04-14 | Central African Republic | I | Selekon,B.,Labouba,I.L.,Gonofio,E.C.,Sem Ouilibona,R.,Simo Tchetgna,H.,Besombes,C.,Feher,M.,Fontanet,A.,Kazanji,M.,Manugerra,J.-C.,Gessain,A.,Nakoune,E.,Berthet,N.,Nkili Meyong,A. |
| Boende_DRC_2008 | KP849469 | 2015-05-13 | 2008-XX-XX | Democratic Republic of the Congo | I | Nakazawa,Y.,Mauldin,M.R.,Emerson,G.L.,Reynolds,M.G.,Lash,R.R.,Gao,J.,Zhao,H.,Li,Y.,Muyembe,J.J.,Kingebeni,P.M.,Wemakoy,O.,Malekani,J.,Karem,K.L.,Damon,I.K.,Carroll,D.S.,Yoshinori,N.,Muyembe,J.-J.,Kingbeni,P.M. |
| Congo_2003_358 | DQ011154 | 2005-09-27 | 2003-XX-XX | Republic of the Congo | I | Likos,A.M.,Sammons,S.A.,Olson,V.A.,Frace,A.M.,Li,Y.,Olsen-Rasmussen,M.,Davidson,W.,Galloway,R.,Khristova,M.L.,Reynolds,M.G.,Zhao,H.,Carroll,D.S.,Curns,A.,Formenty,P.,Esposito,J.J.,Regnery,R.L.,Damon,I.K.,Olsen-Ras |
| 24MPX0037V | PP601196 | 2024-05-02 | 2024-01-05 | Democratic Republic of the Congo | I | Vakaniaki,E.H.,Kaciat,C.,Kinganda - Lusamaki,E.,O'Toole,A.,Wawina -Bokalanga,T.,Mukadi - Bamuleka,D.,Amuri,A.A.,Parker,E.,Muswamba-Kayembe,P.-C.,Makangara - Cigolo,J.-C.,Mulopo - Mukanya,N.,Pukuta - Simbu,E.,M |
| DRC 07-0662 | JX878429 | 2014-01-14 | 2007-09-04 | Democratic Republic of the Congo | I | Kugelman,J.R.,Johnston,S.C.,Mulembakani,P.M.,Kisalu,N.,Lee,M.S.,Koroleva,G.,McCarthy,S.E.,Gestole,M.C.,Wolfe,N.D.,Fair,J.N.,Schneider,B.S.,Wright,L.L.,Huggins,J.,Whitehouse,C.A.,Wemakoy,E.O.,Muyembe-Tamfum,J.J. |
| DRC 07-0354 | JX878425 | 2014-01-14 | 2007-04-20 | Democratic Republic of the Congo | I | Kugelman,J.R.,Johnston,S.C.,Mulembakani,P.M.,Kisalu,N.,Lee,M.S.,Koroleva,G.,McCarthy,S.E.,Gestole,M.C.,Wolfe,N.D.,Fair,J.N.,Schneider,B.S.,Wright,L.L.,Huggins,J.,Whitehouse,C.A.,Wemakoy,E.O.,Muyembe-Tamfum,J.J. |
| DRC 06-0950 | JX878407 | 2014-01-14 | 2006-10-09 | Democratic Republic of the Congo | I | Kugelman,J.R.,Johnston,S.C.,Mulembakani,P.M.,Kisalu,N.,Lee,M.S.,Koroleva,G.,McCarthy,S.E.,Gestole,M.C.,Wolfe,N.D.,Fair,J.N.,Schneider,B.S.,Wright,L.L.,Huggins,J.,Whitehouse,C.A.,Wemakoy,E.O.,Muyembe-Tamfum,J.J. |
| DRC 07-0338 | JX878424 | 2014-01-14 | 2007-03-25 | Democratic Republic of the Congo | I | Kugelman,J.R.,Johnston,S.C.,Mulembakani,P.M.,Kisalu,N.,Lee,M.S.,Koroleva,G.,McCarthy,S.E.,Gestole,M.C.,Wolfe,N.D.,Fair,J.N.,Schneider,B.S.,Wright,L.L.,Huggins,J.,Whitehouse,C.A.,Wemakoy,E.O.,Muyembe-Tamfum,J.J. |
| DRC 07-0337 | JX878423 | 2014-01-14 | 2007-03-25 | Democratic Republic of the Congo | I | Kugelman,J.R.,Johnston,S.C.,Mulembakani,P.M.,Kisalu,N.,Lee,M.S.,Koroleva,G.,McCarthy,S.E.,Gestole,M.C.,Wolfe,N.D.,Fair,J.N.,Schneider,B.S.,Wright,L.L.,Huggins,J.,Whitehouse,C.A.,Wemakoy,E.O.,Muyembe-Tamfum,J.J. |
| 24MPX0194V | PP601206 | 2024-05-02 | 2024-01-20 | Democratic Republic of the Congo | I | Vakaniaki,E.H.,Kaciat,C.,Kinganda - Lusamaki,E.,O'Toole,A.,Wawina -Bokalanga,T.,Mukadi - Bamuleka,D.,Amuri,A.A.,Parker,E.,Muswamba-Kayembe,P.-C.,Makangara - Cigolo,J.-C.,Mulopo - Mukanya,N.,Pukuta - Simbu,E.,M |
| 24MPX0188V | PP601205 | 2024-05-02 | 2024-01-20 | Democratic Republic of the Congo | I | Vakaniaki,E.H.,Kaciat,C.,Kinganda - Lusamaki,E.,O'Toole,A.,Wawina -Bokalanga,T.,Mukadi - Bamuleka,D.,Amuri,A.A.,Parker,E.,Muswamba-Kayembe,P.-C.,Makangara - Cigolo,J.-C.,Mulopo - Mukanya,N.,Pukuta - Simbu,E.,M |
| 23MPX1806V | PP601187 | 2024-05-02 | 2023-12-22 | Democratic Republic of the Congo | I | Vakaniaki,E.H.,Kaciat,C.,Kinganda - Lusamaki,E.,O'Toole,A.,Wawina -Bokalanga,T.,Mukadi - Bamuleka,D.,Amuri,A.A.,Parker,E.,Muswamba-Kayembe,P.-C.,Makangara - Cigolo,J.-C.,Mulopo - Mukanya,N.,Pukuta - Simbu,E.,M |
| 24MPX0009C | PP601189 | 2024-05-02 | 2023-12-18 | Democratic Republic of the Congo | I | Vakaniaki,E.H.,Kaciat,C.,Kinganda - Lusamaki,E.,O'Toole,A.,Wawina -Bokalanga,T.,Mukadi - Bamuleka,D.,Amuri,A.A.,Parker,E.,Muswamba-Kayembe,P.-C.,Makangara - Cigolo,J.-C.,Mulopo - Mukanya,N.,Pukuta - Simbu,E.,M |
| 23MPX1766C | PP601183 | 2024-05-02 | 2023-11-21 | Democratic Republic of the Congo | I | Vakaniaki,E.H.,Kaciat,C.,Kinganda - Lusamaki,E.,O'Toole,A.,Wawina -Bokalanga,T.,Mukadi - Bamuleka,D.,Amuri,A.A.,Parker,E.,Muswamba-Kayembe,P.-C.,Makangara - Cigolo,J.-C.,Mulopo - Mukanya,N.,Pukuta - Simbu,E.,M |
| 24MPX0012C | PP601190 | 2024-05-02 | 2023-12-23 | Democratic Republic of the Congo | I | Vakaniaki,E.H.,Kaciat,C.,Kinganda - Lusamaki,E.,O'Toole,A.,Wawina -Bokalanga,T.,Mukadi - Bamuleka,D.,Amuri,A.A.,Parker,E.,Muswamba-Kayembe,P.-C.,Makangara - Cigolo,J.-C.,Mulopo - Mukanya,N.,Pukuta - Simbu,E.,M |
| MN702453 | MN702453 | 2020-08-29 | 2001-08-11 | Central African Republic | I | Selekon,B.,Labouba,I.L.,Gonofio,E.C.,Sem Ouilibona,R.,Simo Tchetgna,H.,Besombes,C.,Feher,M.,Fontanet,A.,Kazanji,M.,Manugerra,J.-C.,Gessain,A.,Nakoune,E.,Berthet,N.,Nkili Meyong,A. |
| MN702452 | MN702452 | 2020-08-29 | 2010-08-10 | Central African Republic | I | Selekon,B.,Labouba,I.L.,Gonofio,E.C.,Sem Ouilibona,R.,Simo Tchetgna,H.,Besombes,C.,Feher,M.,Fontanet,A.,Kazanji,M.,Manugerra,J.-C.,Gessain,A.,Nakoune,E.,Berthet,N.,Nkili Meyong,A. |
| MN702448 | MN702448 | 2020-08-29 | 2018-03-07 | Central African Republic | I | Selekon,B.,Labouba,I.L.,Gonofio,E.C.,Sem Ouilibona,R.,Simo Tchetgna,H.,Besombes,C.,Feher,M.,Fontanet,A.,Kazanji,M.,Manugerra,J.-C.,Gessain,A.,Nakoune,E.,Berthet,N.,Nkili Meyong,A. |
| MN702447 | MN702447 | 2020-08-29 | 2018-03-20 | Central African Republic | I | Selekon,B.,Labouba,I.L.,Gonofio,E.C.,Sem Ouilibona,R.,Simo Tchetgna,H.,Besombes,C.,Feher,M.,Fontanet,A.,Kazanji,M.,Manugerra,J.-C.,Gessain,A.,Nakoune,E.,Berthet,N.,Nkili Meyong,A. |
| MN702450 | MN702450 | 2020-08-29 | 2016-01-01 | Central African Republic | I | Selekon,B.,Labouba,I.L.,Gonofio,E.C.,Sem Ouilibona,R.,Simo Tchetgna,H.,Besombes,C.,Feher,M.,Fontanet,A.,Kazanji,M.,Manugerra,J.-C.,Gessain,A.,Nakoune,E.,Berthet,N.,Nkili Meyong,A. |
| 24MPX0164V | PP601199 | 2024-05-02 | 2024-01-13 | Democratic Republic of the Congo | I | Vakaniaki,E.H.,Kaciat,C.,Kinganda - Lusamaki,E.,O'Toole,A.,Wawina -Bokalanga,T.,Mukadi - Bamuleka,D.,Amuri,A.A.,Parker,E.,Muswamba-Kayembe,P.-C.,Makangara - Cigolo,J.-C.,Mulopo - Mukanya,N.,Pukuta - Simbu,E.,M |
| 24MPX0166C | PP601200 | 2024-05-02 | 2024-01-13 | Democratic Republic of the Congo | I | Vakaniaki,E.H.,Kaciat,C.,Kinganda - Lusamaki,E.,O'Toole,A.,Wawina -Bokalanga,T.,Mukadi - Bamuleka,D.,Amuri,A.A.,Parker,E.,Muswamba-Kayembe,P.-C.,Makangara - Cigolo,J.-C.,Mulopo - Mukanya,N.,Pukuta - Simbu,E.,M |
| 24MPX0169C | PP601202 | 2024-05-02 | 2024-01-08 | Democratic Republic of the Congo | I | Vakaniaki,E.H.,Kaciat,C.,Kinganda - Lusamaki,E.,O'Toole,A.,Wawina -Bokalanga,T.,Mukadi - Bamuleka,D.,Amuri,A.A.,Parker,E.,Muswamba-Kayembe,P.-C.,Makangara - Cigolo,J.-C.,Mulopo - Mukanya,N.,Pukuta - Simbu,E.,M |
| 24MPX0168V | PP601201 | 2024-05-02 | 2024-01-08 | Democratic Republic of the Congo | I | Vakaniaki,E.H.,Kaciat,C.,Kinganda - Lusamaki,E.,O'Toole,A.,Wawina -Bokalanga,T.,Mukadi - Bamuleka,D.,Amuri,A.A.,Parker,E.,Muswamba-Kayembe,P.-C.,Makangara - Cigolo,J.-C.,Mulopo - Mukanya,N.,Pukuta - Simbu,E.,M |
| 23MPX1786C | PP601185 | 2024-05-02 | 2023-12-18 | Democratic Republic of the Congo | I | Vakaniaki,E.H.,Kaciat,C.,Kinganda - Lusamaki,E.,O'Toole,A.,Wawina -Bokalanga,T.,Mukadi - Bamuleka,D.,Amuri,A.A.,Parker,E.,Muswamba-Kayembe,P.-C.,Makangara - Cigolo,J.-C.,Mulopo - Mukanya,N.,Pukuta - Simbu,E.,M |
| 23MPX1740C | PP601182 | 2024-05-02 | 2023-11-09 | Democratic Republic of the Congo | I | Vakaniaki,E.H.,Kaciat,C.,Kinganda - Lusamaki,E.,O'Toole,A.,Wawina -Bokalanga,T.,Mukadi - Bamuleka,D.,Amuri,A.A.,Parker,E.,Muswamba-Kayembe,P.-C.,Makangara - Cigolo,J.-C.,Mulopo - Mukanya,N.,Pukuta - Simbu,E.,M |
| 23MPX1793V | PP601186 | 2024-05-02 | 2023-12-16 | Democratic Republic of the Congo | I | Vakaniaki,E.H.,Kaciat,C.,Kinganda - Lusamaki,E.,O'Toole,A.,Wawina -Bokalanga,T.,Mukadi - Bamuleka,D.,Amuri,A.A.,Parker,E.,Muswamba-Kayembe,P.-C.,Makangara - Cigolo,J.-C.,Mulopo - Mukanya,N.,Pukuta - Simbu,E.,M |
| 23MPX1769C | PP601184 | 2024-05-02 | 2023-11-25 | Democratic Republic of the Congo | I | Vakaniaki,E.H.,Kaciat,C.,Kinganda - Lusamaki,E.,O'Toole,A.,Wawina -Bokalanga,T.,Mukadi - Bamuleka,D.,Amuri,A.A.,Parker,E.,Muswamba-Kayembe,P.-C.,Makangara - Cigolo,J.-C.,Mulopo - Mukanya,N.,Pukuta - Simbu,E.,M |
| 24MPX0038V | PP601197 | 2024-05-02 | 2024-01-07 | Democratic Republic of the Congo | I | Vakaniaki,E.H.,Kaciat,C.,Kinganda - Lusamaki,E.,O'Toole,A.,Wawina -Bokalanga,T.,Mukadi - Bamuleka,D.,Amuri,A.A.,Parker,E.,Muswamba-Kayembe,P.-C.,Makangara - Cigolo,J.-C.,Mulopo - Mukanya,N.,Pukuta - Simbu,E.,M |
| 24MPX0041V | PP601198 | 2024-05-02 | 2024-01-08 | Democratic Republic of the Congo | I | Vakaniaki,E.H.,Kaciat,C.,Kinganda - Lusamaki,E.,O'Toole,A.,Wawina -Bokalanga,T.,Mukadi - Bamuleka,D.,Amuri,A.A.,Parker,E.,Muswamba-Kayembe,P.-C.,Makangara - Cigolo,J.-C.,Mulopo - Mukanya,N.,Pukuta - Simbu,E.,M |
| 24MPX0014C | PP601191 | 2024-05-02 | 2023-12-24 | Democratic Republic of the Congo | I | Vakaniaki,E.H.,Kaciat,C.,Kinganda - Lusamaki,E.,O'Toole,A.,Wawina -Bokalanga,T.,Mukadi - Bamuleka,D.,Amuri,A.A.,Parker,E.,Muswamba-Kayembe,P.-C.,Makangara - Cigolo,J.-C.,Mulopo - Mukanya,N.,Pukuta - Simbu,E.,M |
| COD/2023/172V | OQ729808 | 2023-04-09 | 2022-XX-XX | Democratic Republic of the Congo | I | Lee,T.Y.,Hwang,Y.-H.,Yun,M.-R.,Kim,Y.-J.,Kim,D. |
| 24MPX0175V | PP601204 | 2024-05-02 | 2024-01-08 | Democratic Republic of the Congo | I | Vakaniaki,E.H.,Kaciat,C.,Kinganda - Lusamaki,E.,O'Toole,A.,Wawina -Bokalanga,T.,Mukadi - Bamuleka,D.,Amuri,A.A.,Parker,E.,Muswamba-Kayembe,P.-C.,Makangara - Cigolo,J.-C.,Mulopo - Mukanya,N.,Pukuta - Simbu,E.,M |
| 24MPX0174C | PP601203 | 2024-05-02 | 2024-01-08 | Democratic Republic of the Congo | I | Vakaniaki,E.H.,Kaciat,C.,Kinganda - Lusamaki,E.,O'Toole,A.,Wawina -Bokalanga,T.,Mukadi - Bamuleka,D.,Amuri,A.A.,Parker,E.,Muswamba-Kayembe,P.-C.,Makangara - Cigolo,J.-C.,Mulopo - Mukanya,N.,Pukuta - Simbu,E.,M |
| 24MPX0025V | PP601194 | 2024-05-02 | 2023-12-30 | Democratic Republic of the Congo | I | Vakaniaki,E.H.,Kaciat,C.,Kinganda - Lusamaki,E.,O'Toole,A.,Wawina -Bokalanga,T.,Mukadi - Bamuleka,D.,Amuri,A.A.,Parker,E.,Muswamba-Kayembe,P.-C.,Makangara - Cigolo,J.-C.,Mulopo - Mukanya,N.,Pukuta - Simbu,E.,M |
| 24MPX0024C | PP601193 | 2024-05-02 | 2023-12-27 | Democratic Republic of the Congo | I | Vakaniaki,E.H.,Kaciat,C.,Kinganda - Lusamaki,E.,O'Toole,A.,Wawina -Bokalanga,T.,Mukadi - Bamuleka,D.,Amuri,A.A.,Parker,E.,Muswamba-Kayembe,P.-C.,Makangara - Cigolo,J.-C.,Mulopo - Mukanya,N.,Pukuta - Simbu,E.,M |
| 24MPX0026V | PP601195 | 2024-05-02 | 2023-12-30 | Democratic Republic of the Congo | I | Vakaniaki,E.H.,Kaciat,C.,Kinganda - Lusamaki,E.,O'Toole,A.,Wawina -Bokalanga,T.,Mukadi - Bamuleka,D.,Amuri,A.A.,Parker,E.,Muswamba-Kayembe,P.-C.,Makangara - Cigolo,J.-C.,Mulopo - Mukanya,N.,Pukuta - Simbu,E.,M |
| 24MPX0018V | PP601192 | 2024-05-02 | 2024-01-02 | Democratic Republic of the Congo | I | Vakaniaki,E.H.,Kaciat,C.,Kinganda - Lusamaki,E.,O'Toole,A.,Wawina -Bokalanga,T.,Mukadi - Bamuleka,D.,Amuri,A.A.,Parker,E.,Muswamba-Kayembe,P.-C.,Makangara - Cigolo,J.-C.,Mulopo - Mukanya,N.,Pukuta - Simbu,E.,M |
| Yambuku_DRC_1985 | KP849471 | 2015-05-13 | 1985-XX-XX | Democratic Republic of the Congo | I | Nakazawa,Y.,Mauldin,M.R.,Emerson,G.L.,Reynolds,M.G.,Lash,R.R.,Gao,J.,Zhao,H.,Li,Y.,Muyembe,J.J.,Kingebeni,P.M.,Wemakoy,O.,Malekani,J.,Karem,K.L.,Damon,I.K.,Carroll,D.S.,Yoshinori,N.,Muyembe,J.-J.,Kingbeni,P.M. |
| DRC 06-1070 | JX878410 | 2014-01-14 | 2006-11-24 | Democratic Republic of the Congo | I | Kugelman,J.R.,Johnston,S.C.,Mulembakani,P.M.,Kisalu,N.,Lee,M.S.,Koroleva,G.,McCarthy,S.E.,Gestole,M.C.,Wolfe,N.D.,Fair,J.N.,Schneider,B.S.,Wright,L.L.,Huggins,J.,Whitehouse,C.A.,Wemakoy,E.O.,Muyembe-Tamfum,J.J. |
| DRC 06-1076 | JX878412 | 2014-01-14 | 2006-11-30 | Democratic Republic of the Congo | I | Kugelman,J.R.,Johnston,S.C.,Mulembakani,P.M.,Kisalu,N.,Lee,M.S.,Koroleva,G.,McCarthy,S.E.,Gestole,M.C.,Wolfe,N.D.,Fair,J.N.,Schneider,B.S.,Wright,L.L.,Huggins,J.,Whitehouse,C.A.,Wemakoy,E.O.,Muyembe-Tamfum,J.J. |
| DRC 06-1075 | JX878411 | 2014-01-14 | 2006-11-30 | Democratic Republic of the Congo | I | Kugelman,J.R.,Johnston,S.C.,Mulembakani,P.M.,Kisalu,N.,Lee,M.S.,Koroleva,G.,McCarthy,S.E.,Gestole,M.C.,Wolfe,N.D.,Fair,J.N.,Schneider,B.S.,Wright,L.L.,Huggins,J.,Whitehouse,C.A.,Wemakoy,E.O.,Muyembe-Tamfum,J.J. |
| DRC 06-0999 | JX878409 | 2014-01-14 | 2006-11-09 | Democratic Republic of the Congo | I | Kugelman,J.R.,Johnston,S.C.,Mulembakani,P.M.,Kisalu,N.,Lee,M.S.,Koroleva,G.,McCarthy,S.E.,Gestole,M.C.,Wolfe,N.D.,Fair,J.N.,Schneider,B.S.,Wright,L.L.,Huggins,J.,Whitehouse,C.A.,Wemakoy,E.O.,Muyembe-Tamfum,J.J. |
| DRC 07-0287 | JX878422 | 2014-01-14 | 2007-03-20 | Democratic Republic of the Congo | I | Kugelman,J.R.,Johnston,S.C.,Mulembakani,P.M.,Kisalu,N.,Lee,M.S.,Koroleva,G.,McCarthy,S.E.,Gestole,M.C.,Wolfe,N.D.,Fair,J.N.,Schneider,B.S.,Wright,L.L.,Huggins,J.,Whitehouse,C.A.,Wemakoy,E.O.,Muyembe-Tamfum,J.J. |
| DRC 07-0514 | JX878428 | 2014-01-14 | 2007-06-30 | Democratic Republic of the Congo | I | Kugelman,J.R.,Johnston,S.C.,Mulembakani,P.M.,Kisalu,N.,Lee,M.S.,Koroleva,G.,McCarthy,S.E.,Gestole,M.C.,Wolfe,N.D.,Fair,J.N.,Schneider,B.S.,Wright,L.L.,Huggins,J.,Whitehouse,C.A.,Wemakoy,E.O.,Muyembe-Tamfum,J.J. |
| DRC 07-0046 | JX878414 | 2014-01-14 | 2006-12-14 | Democratic Republic of the Congo | I | Kugelman,J.R.,Johnston,S.C.,Mulembakani,P.M.,Kisalu,N.,Lee,M.S.,Koroleva,G.,McCarthy,S.E.,Gestole,M.C.,Wolfe,N.D.,Fair,J.N.,Schneider,B.S.,Wright,L.L.,Huggins,J.,Whitehouse,C.A.,Wemakoy,E.O.,Muyembe-Tamfum,J.J. |
| DRC 07-0480 | JX878427 | 2014-01-14 | 2007-05-25 | Democratic Republic of the Congo | I | Kugelman,J.R.,Johnston,S.C.,Mulembakani,P.M.,Kisalu,N.,Lee,M.S.,Koroleva,G.,McCarthy,S.E.,Gestole,M.C.,Wolfe,N.D.,Fair,J.N.,Schneider,B.S.,Wright,L.L.,Huggins,J.,Whitehouse,C.A.,Wemakoy,E.O.,Muyembe-Tamfum,J.J. |
| DRC 07-0286 | JX878421 | 2014-01-14 | 2007-03-22 | Democratic Republic of the Congo | I | Kugelman,J.R.,Johnston,S.C.,Mulembakani,P.M.,Kisalu,N.,Lee,M.S.,Koroleva,G.,McCarthy,S.E.,Gestole,M.C.,Wolfe,N.D.,Fair,J.N.,Schneider,B.S.,Wright,L.L.,Huggins,J.,Whitehouse,C.A.,Wemakoy,E.O.,Muyembe-Tamfum,J.J. |
| DRC 07-0093 | JX878416 | 2014-01-14 | 2006-12-26 | Democratic Republic of the Congo | I | Kugelman,J.R.,Johnston,S.C.,Mulembakani,P.M.,Kisalu,N.,Lee,M.S.,Koroleva,G.,McCarthy,S.E.,Gestole,M.C.,Wolfe,N.D.,Fair,J.N.,Schneider,B.S.,Wright,L.L.,Huggins,J.,Whitehouse,C.A.,Wemakoy,E.O.,Muyembe-Tamfum,J.J. |
| DRC 07-0092 | JX878415 | 2014-01-14 | 2006-12-26 | Democratic Republic of the Congo | I | Kugelman,J.R.,Johnston,S.C.,Mulembakani,P.M.,Kisalu,N.,Lee,M.S.,Koroleva,G.,McCarthy,S.E.,Gestole,M.C.,Wolfe,N.D.,Fair,J.N.,Schneider,B.S.,Wright,L.L.,Huggins,J.,Whitehouse,C.A.,Wemakoy,E.O.,Muyembe-Tamfum,J.J. |
| DRC 07-0045 | JX878413 | 2014-01-14 | 2006-12-14 | Democratic Republic of the Congo | I | Kugelman,J.R.,Johnston,S.C.,Mulembakani,P.M.,Kisalu,N.,Lee,M.S.,Koroleva,G.,McCarthy,S.E.,Gestole,M.C.,Wolfe,N.D.,Fair,J.N.,Schneider,B.S.,Wright,L.L.,Huggins,J.,Whitehouse,C.A.,Wemakoy,E.O.,Muyembe-Tamfum,J.J. |
| BNITM-Gabon1988 | OP498046 | 2022-09-27 | 1988-XX-XX | Gabon | I | Emmerich,P.,Bialonski,A.,Tomazatos,A.,Cadar,D. |
| KJ642619 | KJ642619 | 2015-05-11 | 1988-XX-XX | Gabon | I | Nakazawa,Y.,Mauldin,M.R.,Emerson,G.L.,Reynolds,M.G.,Lash,R.R.,Gao,J.,Zhao,H.,Li,Y.,Muyembe,J.J.,Kingebeni,P.M.,Wemakoy,O.,Malekani,J.,Karem,K.L.,Damon,I.K.,Carroll,D.S.,Lil,Y.,Weil,R.M.,Muyembe,J.-J.,Kingbeni,P.M |
| KJ642618 | KJ642618 | 2015-05-11 | 1990-XX-XX | Cameroon | I | Nakazawa,Y.,Mauldin,M.R.,Emerson,G.L.,Reynolds,M.G.,Lash,R.R.,Gao,J.,Zhao,H.,Li,Y.,Muyembe,J.J.,Kingebeni,P.M.,Wemakoy,O.,Malekani,J.,Karem,K.L.,Damon,I.K.,Carroll,D.S.,Lil,Y.,Weil,R.M.,Muyembe,J.-J.,Kingbeni,P.M |
| CMR_2016_Chimp | OR943698 | 2023-12-25 | 2016-08-XX | Cameroon | I | Brien,S.C.,LeBreton,M.,Doty,J.B.,Mauldin,M.R.,Morgan,C.N.,Pieracci,E.G.,Ritter,J.M.,Matheny,A.,Wilkins,K.,Tafon,B.G.,Tamoufe,U.,Missoup,A.D.,Nwobegahay,J.,Takuo,J.M.,Nkom,F.,Mouiche,M.M.,Feussom,J.M.,Wade,A.,Mc |
| KJ642612 | KJ642612 | 2015-05-11 | 1986-XX-XX | Democratic Republic of the Congo | I | Nakazawa,Y.,Mauldin,M.R.,Emerson,G.L.,Reynolds,M.G.,Lash,R.R.,Gao,J.,Zhao,H.,Li,Y.,Muyembe,J.J.,Kingebeni,P.M.,Wemakoy,O.,Malekani,J.,Karem,K.L.,Damon,I.K.,Carroll,D.S.,Lil,Y.,Weil,R.M.,Muyembe,J.-J.,Kingbeni,P.M |
| DRC 07-0283 | JX878420 | 2014-01-14 | 2007-02-26 | Democratic Republic of the Congo | I | Kugelman,J.R.,Johnston,S.C.,Mulembakani,P.M.,Kisalu,N.,Lee,M.S.,Koroleva,G.,McCarthy,S.E.,Gestole,M.C.,Wolfe,N.D.,Fair,J.N.,Schneider,B.S.,Wright,L.L.,Huggins,J.,Whitehouse,C.A.,Wemakoy,E.O.,Muyembe-Tamfum,J.J. |
| DRC 07-0275 | JX878419 | 2014-01-14 | 2007-02-10 | Democratic Republic of the Congo | I | Kugelman,J.R.,Johnston,S.C.,Mulembakani,P.M.,Kisalu,N.,Lee,M.S.,Koroleva,G.,McCarthy,S.E.,Gestole,M.C.,Wolfe,N.D.,Fair,J.N.,Schneider,B.S.,Wright,L.L.,Huggins,J.,Whitehouse,C.A.,Wemakoy,E.O.,Muyembe-Tamfum,J.J. |
| DRC 07-0450 | JX878426 | 2014-01-14 | 2007-05-27 | Democratic Republic of the Congo | I | Kugelman,J.R.,Johnston,S.C.,Mulembakani,P.M.,Kisalu,N.,Lee,M.S.,Koroleva,G.,McCarthy,S.E.,Gestole,M.C.,Wolfe,N.D.,Fair,J.N.,Schneider,B.S.,Wright,L.L.,Huggins,J.,Whitehouse,C.A.,Wemakoy,E.O.,Muyembe-Tamfum,J.J. |
| DRC 06-0970 | JX878408 | 2014-01-14 | 2006-10-31 | Democratic Republic of the Congo | I | Kugelman,J.R.,Johnston,S.C.,Mulembakani,P.M.,Kisalu,N.,Lee,M.S.,Koroleva,G.,McCarthy,S.E.,Gestole,M.C.,Wolfe,N.D.,Fair,J.N.,Schneider,B.S.,Wright,L.L.,Huggins,J.,Whitehouse,C.A.,Wemakoy,E.O.,Muyembe-Tamfum,J.J. |
| DRC 07-0120 | JX878418 | 2014-01-14 | 2007-01-02 | Democratic Republic of the Congo | I | Kugelman,J.R.,Johnston,S.C.,Mulembakani,P.M.,Kisalu,N.,Lee,M.S.,Koroleva,G.,McCarthy,S.E.,Gestole,M.C.,Wolfe,N.D.,Fair,J.N.,Schneider,B.S.,Wright,L.L.,Huggins,J.,Whitehouse,C.A.,Wemakoy,E.O.,Muyembe-Tamfum,J.J. |
| NC_003310 | NC_003310 | 2001-12-12 | 1996-XX-XX | Democratic Republic of the Congo | I | Senkevich,T.G.,Yutin,N.,Wolf,Y.I.,Koonin,E.V.,Moss,B.,Zmasek,C.M.,Knipe,D.M.,Pellett,P.E.,Scheuermann,R.H.,Shchelkunov,S.N.,Totmenin,A.V.,Babkin,I.V.,Safronov,P.F.,Ryazankina,O.I.,Petrov,N.A.,Gutorov,V.V.,Uvarova,E.A.,M |
| Zaire-96-I-16 | AF380138 | 2001-12-12 | 1996-XX-XX | Democratic Republic of the Congo | I | Shchelkunov,S.N.,Totmenin,A.V.,Babkin,I.V.,Safronov,P.F.,Ryazankina,O.I.,Petrov,N.A.,Gutorov,V.V.,Uvarova,E.A.,Mikheev,M.V.,Sisler,J.R.,Esposito,J.J.,Jahrling,P.B.,Moss,B.,Sandakhchiev,L.S. |
| DRC 07-0104 | JX878417 | 2014-01-14 | 2006-12-14 | Democratic Republic of the Congo | I | Kugelman,J.R.,Johnston,S.C.,Mulembakani,P.M.,Kisalu,N.,Lee,M.S.,Koroleva,G.,McCarthy,S.E.,Gestole,M.C.,Wolfe,N.D.,Fair,J.N.,Schneider,B.S.,Wright,L.L.,Huggins,J.,Whitehouse,C.A.,Wemakoy,E.O.,Muyembe-Tamfum,J.J. |
| RDC-NKV-GOM-MPO | PP601227 | 2024-05-02 | 2023-10-05 | Democratic Republic of the Congo | I | Vakaniaki,E.H.,Kaciat,C.,Kinganda - Lusamaki,E.,O'Toole,A.,Wawina -Bokalanga,T.,Mukadi - Bamuleka,D.,Amuri,A.A.,Parker,E.,Muswamba-Kayembe,P.-C.,Makangara - Cigolo,J.-C.,Mulopo - Mukanya,N.,Pukuta - Simbu,E.,M |
| RDC-NKV-GOM-MPO | PP601228 | 2024-05-02 | 2023-10-13 | Democratic Republic of the Congo | I | Vakaniaki,E.H.,Kaciat,C.,Kinganda - Lusamaki,E.,O'Toole,A.,Wawina -Bokalanga,T.,Mukadi - Bamuleka,D.,Amuri,A.A.,Parker,E.,Muswamba-Kayembe,P.-C.,Makangara - Cigolo,J.-C.,Mulopo - Mukanya,N.,Pukuta - Simbu,E.,M |
| 24MPX0223C | PP601218 | 2024-05-02 | 2023-12-26 | Democratic Republic of the Congo | I | Vakaniaki,E.H.,Kaciat,C.,Kinganda - Lusamaki,E.,O'Toole,A.,Wawina -Bokalanga,T.,Mukadi - Bamuleka,D.,Amuri,A.A.,Parker,E.,Muswamba-Kayembe,P.-C.,Makangara - Cigolo,J.-C.,Mulopo - Mukanya,N.,Pukuta - Simbu,E.,M |
| 24MPX0226V | PP601220 | 2024-05-02 | 2023-12-26 | Democratic Republic of the Congo | I | Vakaniaki,E.H.,Kaciat,C.,Kinganda - Lusamaki,E.,O'Toole,A.,Wawina -Bokalanga,T.,Mukadi - Bamuleka,D.,Amuri,A.A.,Parker,E.,Muswamba-Kayembe,P.-C.,Makangara - Cigolo,J.-C.,Mulopo - Mukanya,N.,Pukuta - Simbu,E.,M |
| 24MPX0214V | PP601214 | 2024-05-02 | 2024-01-23 | Democratic Republic of the Congo | I | Vakaniaki,E.H.,Kaciat,C.,Kinganda - Lusamaki,E.,O'Toole,A.,Wawina -Bokalanga,T.,Mukadi - Bamuleka,D.,Amuri,A.A.,Parker,E.,Muswamba-Kayembe,P.-C.,Makangara - Cigolo,J.-C.,Mulopo - Mukanya,N.,Pukuta - Simbu,E.,M |
| 24MPX0203V | PP601209 | 2024-05-02 | 2024-01-23 | Democratic Republic of the Congo | I | Vakaniaki,E.H.,Kaciat,C.,Kinganda - Lusamaki,E.,O'Toole,A.,Wawina -Bokalanga,T.,Mukadi - Bamuleka,D.,Amuri,A.A.,Parker,E.,Muswamba-Kayembe,P.-C.,Makangara - Cigolo,J.-C.,Mulopo - Mukanya,N.,Pukuta - Simbu,E.,M |
| 24MPX0240V | PP601225 | 2024-05-02 | 2024-01-14 | Democratic Republic of the Congo | I | Vakaniaki,E.H.,Kaciat,C.,Kinganda - Lusamaki,E.,O'Toole,A.,Wawina -Bokalanga,T.,Mukadi - Bamuleka,D.,Amuri,A.A.,Parker,E.,Muswamba-Kayembe,P.-C.,Makangara - Cigolo,J.-C.,Mulopo - Mukanya,N.,Pukuta - Simbu,E.,M |
| 24MPX0224C | PP601219 | 2024-05-02 | 2023-12-25 | Democratic Republic of the Congo | I | Vakaniaki,E.H.,Kaciat,C.,Kinganda - Lusamaki,E.,O'Toole,A.,Wawina -Bokalanga,T.,Mukadi - Bamuleka,D.,Amuri,A.A.,Parker,E.,Muswamba-Kayembe,P.-C.,Makangara - Cigolo,J.-C.,Mulopo - Mukanya,N.,Pukuta - Simbu,E.,M |
| 24MPX0231V | PP601223 | 2024-05-02 | 2023-12-21 | Democratic Republic of the Congo | I | Vakaniaki,E.H.,Kaciat,C.,Kinganda - Lusamaki,E.,O'Toole,A.,Wawina -Bokalanga,T.,Mukadi - Bamuleka,D.,Amuri,A.A.,Parker,E.,Muswamba-Kayembe,P.-C.,Makangara - Cigolo,J.-C.,Mulopo - Mukanya,N.,Pukuta - Simbu,E.,M |
| 24MPX0230C | PP601222 | 2024-05-02 | 2023-12-31 | Democratic Republic of the Congo | I | Vakaniaki,E.H.,Kaciat,C.,Kinganda - Lusamaki,E.,O'Toole,A.,Wawina -Bokalanga,T.,Mukadi - Bamuleka,D.,Amuri,A.A.,Parker,E.,Muswamba-Kayembe,P.-C.,Makangara - Cigolo,J.-C.,Mulopo - Mukanya,N.,Pukuta - Simbu,E.,M |
| 24MPX0209V | PP601213 | 2024-05-02 | 2024-01-19 | Democratic Republic of the Congo | I | Vakaniaki,E.H.,Kaciat,C.,Kinganda - Lusamaki,E.,O'Toole,A.,Wawina -Bokalanga,T.,Mukadi - Bamuleka,D.,Amuri,A.A.,Parker,E.,Muswamba-Kayembe,P.-C.,Makangara - Cigolo,J.-C.,Mulopo - Mukanya,N.,Pukuta - Simbu,E.,M |
| 24MPX0221V | PP601217 | 2024-05-02 | 2024-01-26 | Democratic Republic of the Congo | I | Vakaniaki,E.H.,Kaciat,C.,Kinganda - Lusamaki,E.,O'Toole,A.,Wawina -Bokalanga,T.,Mukadi - Bamuleka,D.,Amuri,A.A.,Parker,E.,Muswamba-Kayembe,P.-C.,Makangara - Cigolo,J.-C.,Mulopo - Mukanya,N.,Pukuta - Simbu,E.,M |
| 24MPX0228V | PP601221 | 2024-05-02 | 2023-12-27 | Democratic Republic of the Congo | I | Vakaniaki,E.H.,Kaciat,C.,Kinganda - Lusamaki,E.,O'Toole,A.,Wawina -Bokalanga,T.,Mukadi - Bamuleka,D.,Amuri,A.A.,Parker,E.,Muswamba-Kayembe,P.-C.,Makangara - Cigolo,J.-C.,Mulopo - Mukanya,N.,Pukuta - Simbu,E.,M |
| 24MPX0201V | PP601208 | 2024-05-02 | 2024-01-23 | Democratic Republic of the Congo | I | Vakaniaki,E.H.,Kaciat,C.,Kinganda - Lusamaki,E.,O'Toole,A.,Wawina -Bokalanga,T.,Mukadi - Bamuleka,D.,Amuri,A.A.,Parker,E.,Muswamba-Kayembe,P.-C.,Makangara - Cigolo,J.-C.,Mulopo - Mukanya,N.,Pukuta - Simbu,E.,M |
| 24MPX0239V | PP601224 | 2024-05-02 | 2024-01-10 | Democratic Republic of the Congo | I | Vakaniaki,E.H.,Kaciat,C.,Kinganda - Lusamaki,E.,O'Toole,A.,Wawina -Bokalanga,T.,Mukadi - Bamuleka,D.,Amuri,A.A.,Parker,E.,Muswamba-Kayembe,P.-C.,Makangara - Cigolo,J.-C.,Mulopo - Mukanya,N.,Pukuta - Simbu,E.,M |
| 24MPX0220V | PP601216 | 2024-05-02 | 2024-01-26 | Democratic Republic of the Congo | I | Vakaniaki,E.H.,Kaciat,C.,Kinganda - Lusamaki,E.,O'Toole,A.,Wawina -Bokalanga,T.,Mukadi - Bamuleka,D.,Amuri,A.A.,Parker,E.,Muswamba-Kayembe,P.-C.,Makangara - Cigolo,J.-C.,Mulopo - Mukanya,N.,Pukuta - Simbu,E.,M |
| 24MPX0205V | PP601210 | 2024-05-02 | 2024-01-28 | Democratic Republic of the Congo | I | Vakaniaki,E.H.,Kaciat,C.,Kinganda - Lusamaki,E.,O'Toole,A.,Wawina -Bokalanga,T.,Mukadi - Bamuleka,D.,Amuri,A.A.,Parker,E.,Muswamba-Kayembe,P.-C.,Makangara - Cigolo,J.-C.,Mulopo - Mukanya,N.,Pukuta - Simbu,E.,M |
| 24MPX0242V | PP601226 | 2024-05-02 | 2024-01-15 | Democratic Republic of the Congo | I | Vakaniaki,E.H.,Kaciat,C.,Kinganda - Lusamaki,E.,O'Toole,A.,Wawina -Bokalanga,T.,Mukadi - Bamuleka,D.,Amuri,A.A.,Parker,E.,Muswamba-Kayembe,P.-C.,Makangara - Cigolo,J.-C.,Mulopo - Mukanya,N.,Pukuta - Simbu,E.,M |
| 24MPX0207Or | PP601212 | 2024-05-02 | 2024-01-23 | Democratic Republic of the Congo | I | Vakaniaki,E.H.,Kaciat,C.,Kinganda - Lusamaki,E.,O'Toole,A.,Wawina -Bokalanga,T.,Mukadi - Bamuleka,D.,Amuri,A.A.,Parker,E.,Muswamba-Kayembe,P.-C.,Makangara - Cigolo,J.-C.,Mulopo - Mukanya,N.,Pukuta - Simbu,E.,M |
| 24MPX0198V | PP601207 | 2024-05-02 | 2024-01-23 | Democratic Republic of the Congo | I | Vakaniaki,E.H.,Kaciat,C.,Kinganda - Lusamaki,E.,O'Toole,A.,Wawina -Bokalanga,T.,Mukadi - Bamuleka,D.,Amuri,A.A.,Parker,E.,Muswamba-Kayembe,P.-C.,Makangara - Cigolo,J.-C.,Mulopo - Mukanya,N.,Pukuta - Simbu,E.,M |
| 24MPX0217V | PP601215 | 2024-05-02 | 2024-01-23 | Democratic Republic of the Congo | I | Vakaniaki,E.H.,Kaciat,C.,Kinganda - Lusamaki,E.,O'Toole,A.,Wawina -Bokalanga,T.,Mukadi - Bamuleka,D.,Amuri,A.A.,Parker,E.,Muswamba-Kayembe,P.-C.,Makangara - Cigolo,J.-C.,Mulopo - Mukanya,N.,Pukuta - Simbu,E.,M |
| 24MPX0206V | PP601211 | 2024-05-02 | 2024-01-23 | Democratic Republic of the Congo | I | Vakaniaki,E.H.,Kaciat,C.,Kinganda - Lusamaki,E.,O'Toole,A.,Wawina -Bokalanga,T.,Mukadi - Bamuleka,D.,Amuri,A.A.,Parker,E.,Muswamba-Kayembe,P.-C.,Makangara - Cigolo,J.-C.,Mulopo - Mukanya,N.,Pukuta - Simbu,E.,M |
| USA2003_099_Gambia | MT903346 | 2020-09-15 | 2003-XX-XX | USA | IIa | Mauldin,M.R.,McCollum,A.M.,Nakazawa,Y.J.,Mandra,A.,Whitehouse,E.R.,Davidson,W.,Zhao,H.,Gao,J.,Li,Y.,Doty,J.,Yinka-Ogunleye,A.,Akinpelu,A.,Aruna,O.,Naidoo,D.,Lewandowski,K.,Afrough,B.,Graham,V.,Aarons,E.,Hewsor |
| DQ011153 | DQ011153 | 2005-09-27 | 2003-XX-XX | USA | IIa | Likos,A.M.,Sammons,S.A.,Olson,V.A.,Frace,A.M.,Li,Y.,Olsen-Rasmussen,M.,Davidson,W.,Galloway,R.,Khristova,M.L.,Reynolds,M.G.,Zhao,H.,Carroll,D.S.,Curns,A.,Formenty,P.,Esposito,J.J.,Regnery,R.L.,Damon,I.K.,Olsen-Ras |
| KJ642616 | KJ642616 | 2015-05-11 | 1968-XX-XX | France | IIa | Nakazawa,Y.,Mauldin,M.R.,Emerson,G.L.,Reynolds,M.G.,Lash,R.R.,Gao,J.,Zhao,H.,Li,Y.,Muyembe,J.J.,Kingebeni,P.M.,Wemakoy,O.,Malekani,J.,Karem,K.L.,Damon,I.K.,Carroll,D.S.,Lil,Y.,Weil,R.M.,Muyembe,J.-J.,Kingbeni,P.M |
| KJ642614 | KJ642614 | 2015-05-11 | 1965-XX-XX | Netherlands | IIa | Nakazawa,Y.,Mauldin,M.R.,Emerson,G.L.,Reynolds,M.G.,Lash,R.R.,Gao,J.,Zhao,H.,Li,Y.,Muyembe,J.J.,Kingebeni,P.M.,Wemakoy,O.,Malekani,J.,Karem,K.L.,Damon,I.K.,Carroll,D.S.,Lil,Y.,Weil,R.M.,Muyembe,J.-J.,Kingbeni,P.M |
| AY753185 | AY753185 | 2005-09-07 | 1958-XX-XX |  | IIa | Chen,N.,Li,G.,Liszewski,M.K.,Atkinson,J.P.,Jahrling,P.B.,Feng,Z.,Schriewer,J.,Buck,C.,Wang,C.,Lefkowitz,E.J.,Esposito,J.J.,Harms,T.,Damon,I.K.,Roper,R.L.,Upton,C.,Buller,R.M.,Buller,R.M.L. |
| AY603973 | AY603973 | 2004-12-30 | 1961-XX-XX |  | IIa | Chen,N.,Li,G.,Liszewski,M.K.,Atkinson,J.P.,Jahrling,P.B.,Feng,Z.,Schriewer,J.,Buck,C.,Wang,C.,Lefkowitz,E.J.,Esposito,J.J.,Harms,T.,Damon,I.K.,Roper,R.L.,Upton,C.,Buller,R.M.,Buller,R.M.L. |
| Sierra Leone | AY741551 | 2005-09-07 | 1970-XX-XX | Sierra Leone | IIa | Chen,N.,Li,G.,Liszewski,M.K.,Atkinson,J.P.,Jahrling,P.B.,Feng,Z.,Schriewer,J.,Buck,C.,Wang,C.,Lefkowitz,E.J.,Esposito,J.J.,Harms,T.,Damon,I.K.,Roper,R.L.,Upton,C.,Buller,R.M.,Buller,R.M.L. |
| Cote d'Ivoire_1971 | KP849470 | 2015-05-13 | 1971-XX-XX | Côte d'Ivoire | IIa | Nakazawa,Y.,Mauldin,M.R.,Emerson,G.L.,Reynolds,M.G.,Lash,R.R.,Gao,J.,Zhao,H.,Li,Y.,Muyembe,J.J.,Kingebeni,P.M.,Wemakoy,O.,Malekani,J.,Karem,K.L.,Damon,I.K.,Carroll,D.S.,Yoshinori,N.,Muyembe,J.-J.,Kingbeni,P.M. |
| DQ011156 | DQ011156 | 2005-09-27 | 1970-XX-XX | Liberia | IIa | Likos,A.M.,Sammons,S.A.,Olson,V.A.,Frace,A.M.,Li,Y.,Olsen-Rasmussen,M.,Davidson,W.,Galloway,R.,Khristova,M.L.,Reynolds,M.G.,Zhao,H.,Carroll,D.S.,Curns,A.,Formenty,P.,Esposito,J.J.,Regnery,R.L.,Damon,I.K.,Olsen-Ras |
| KJ136820 | KJ136820 | 2014-03-30 | 2012-03-XX | Côte d'Ivoire | IIa | Radonic,A.,Metzger,S.,Dabrowski,P.W.,Couacy-Hymann,E.,Schuenadel,L.,Kurth,A.,Matz-Rensing,K.,Boesch,C.,Leendertz,F.H.,Nitsche,A. |
| TNP_2017_North_Surp | MN346697 | 2020-03-17 | 2017-04-14 | Côte d'Ivoire | IIa | Patrono,L.V.,Pleh,K.,Samuni,L.,Ulrich,M.,Roethemeier,C.,Sachse,A.,Muschter,S.,Nitsche,A.,Couacy-Hymann,E.,Boesch,C.,Wittig,R.M.,Calvignac-Spencer,S.,Leendertz,F.H. |
| TNP_2017_North_Sarc | MN346694 | 2020-03-17 | 2017-03-28 | Côte d'Ivoire | IIa | Patrono,L.V.,Pleh,K.,Samuni,L.,Ulrich,M.,Roethemeier,C.,Sachse,A.,Muschter,S.,Nitsche,A.,Couacy-Hymann,E.,Boesch,C.,Wittig,R.M.,Calvignac-Spencer,S.,Leendertz,F.H. |
| KJ642615 | KJ642615 | 2015-05-11 | 1978-XX-XX | Nigeria | IIb | Nakazawa,Y.,Mauldin,M.R.,Emerson,G.L.,Reynolds,M.G.,Lash,R.R.,Gao,J.,Zhao,H.,Li,Y.,Muyembe,J.J.,Kingebeni,P.M.,Wemakoy,O.,Malekani,J.,Karem,K.L.,Damon,I.K.,Carroll,D.S.,Lil,Y.,Weil,R.M.,Muyembe,J.-J.,Kingbeni,P.M |
| 21V-04877 | PP860031 | 2024-07-09 | 2021-07-22 | Cameroon | IIb | Djuicy,D.D.,Omah,I.F.,Parker,E.,Tomkins-Tinch,C.H.,Otieno,J.R.,Yifomnjou,M.H.M.,Essengue,L.L.M.,Ayinla,A.O.,Sijuwola,A.E.,Ahmed,M.I.,Ope-ewe,O.O.,Ogunsanya,O.A.,Olono,A.,Eromon,P.,Yonga,M.G.W.,Essima,G.D.,Touoy |
| 22V-07968 | PP860027 | 2024-07-09 | 2022-10-13 | Cameroon | IIb | Djuicy,D.D.,Omah,I.F.,Parker,E.,Tomkins-Tinch,C.H.,Otieno,J.R.,Yifomnjou,M.H.M.,Essengue,L.L.M.,Ayinla,A.O.,Sijuwola,A.E.,Ahmed,M.I.,Ope-ewe,O.O.,Ogunsanya,O.A.,Olono,A.,Eromon,P.,Yonga,M.G.W.,Essima,G.D.,Touoy |
| KJ642617 | KJ642617 | 2015-05-11 | 1971-XX-XX | Nigeria | IIb | Nakazawa,Y.,Mauldin,M.R.,Emerson,G.L.,Reynolds,M.G.,Lash,R.R.,Gao,J.,Zhao,H.,Li,Y.,Muyembe,J.J.,Kingebeni,P.M.,Wemakoy,O.,Malekani,J.,Karem,K.L.,Damon,I.K.,Carroll,D.S.,Lil,Y.,Weil,R.M.,Muyembe,J.-J.,Kingbeni,P.M |
| 046/19 | OP612684 | 2022-10-11 | 2019-05-XX | Nigeria | IIb | Ndodo,N.,Ashcroft,J.,Lewandowski,K.,Yinka-Ogunleye,A.,Chukwu,C.,Ahmad,A.,King,D.,Akinpelu,A.,Maluquer de Motes,C.,Ribeca,P.,Summer,R.P.,Rambaut,A.,Chester,M.,Maishman,T.,Babatunde,O.,Mba,N.,Aruna,O.,Pullan,S |
| 191/20 | OP612691 | 2022-10-11 | 2020-01-XX | Nigeria | IIb | Ndodo,N.,Ashcroft,J.,Lewandowski,K.,Yinka-Ogunleye,A.,Chukwu,C.,Ahmad,A.,King,D.,Akinpelu,A.,Maluquer de Motes,C.,Ribeca,P.,Summer,R.P.,Rambaut,A.,Chester,M.,Maishman,T.,Babatunde,O.,Mba,N.,Aruna,O.,Pullan,S |
| TRM328 | PP853022 | 2024-07-09 | 2022-09-16 | Nigeria | IIb | Parker,E.,Omah,I.F.,Varilly,P.,Magee,A.,Ayinla,A.O.,Sijuwola,A.E.,Ahmed,M.I.,Ope-ewe,O.O.,Ogunsanya,O.A.,Olono,A.,Eromon,P.,Tomkins-Tinch,C.H.,Otieno,J.R.,Akanbi,O.,Egwuenu,A.,Ehiakhamen,O.,Chukwu,C.,Suleiman,K. |
| 111/19 | OP612689 | 2022-10-11 | 2019-12-17 | Nigeria | IIb | Ndodo,N.,Ashcroft,J.,Lewandowski,K.,Yinka-Ogunleye,A.,Chukwu,C.,Ahmad,A.,King,D.,Akinpelu,A.,Maluquer de Motes,C.,Ribeca,P.,Summer,R.P.,Rambaut,A.,Chester,M.,Maishman,T.,Babatunde,O.,Mba,N.,Aruna,O.,Pullan,S |

| | | | | | | |
|---|---|---|---|---|---|---|
| **USA_2021_TX** | ON676707 | 2022-06-03 | 2021-07-XX | USA | IIb | Gigante,C.M.,Stringer,J.,Seabolt,M.H.,Wilkins,K.,McCollum,A.,Hutson,C.,Davidson,W.,Rao,A.,Schulte,J.,Li,Y. |
| **VSP208** | PP852966 | 2024-07-09 | 2023-03-15 | Nigeria | IIb | Parker,E.,Omah,I.F.,Varilly,P.,Magee,A.,Ayinla,A.O.,Sijuwola,A.E.,Ahmed,M.I.,Ope-ewe,O.O.,Ogunsanya,O.A.,Olono,A.,Eromon,P.,Tomkins-Tinch,C.H.,Otieno,J.R.,Akanbi,O.,Egwuenu,A.,Ehiakhamen,O.,Chukwu,C.,Suleiman,K. |
| **VSP192** | PP852950 | 2024-07-09 | 2023-01-08 | Nigeria | IIb | Parker,E.,Omah,I.F.,Varilly,P.,Magee,A.,Ayinla,A.O.,Sijuwola,A.E.,Ahmed,M.I.,Ope-ewe,O.O.,Ogunsanya,O.A.,Olono,A.,Eromon,P.,Tomkins-Tinch,C.H.,Otieno,J.R.,Akanbi,O.,Egwuenu,A.,Ehiakhamen,O.,Chukwu,C.,Suleiman,K. |
| **M5320_M15_Bayelsa** | MT903341 | 2020-09-15 | 2018-08-14 | Nigeria | IIb | Mauldin,M.R.,McCollum,A.M.,Nakazawa,Y.J.,Mandra,A.,Whitehouse,E.R.,Davidson,W.,Zhao,H.,Gao,J.,Li,Y.,Doty,J.,Yinka-Ogunleye,A.,Akinpelu,A.,Aruna,O.,Naidoo,D.,Lewandowski,K.,Afrough,B.,Graham,V.,Aarons,E.,Hewsor |
| **MN648051** | MN648051 | 2019-12-07 | 2018-10-04 | Israel | IIb | Cohen Gihon,I.,Israeli,O.,Shifman,O.,Erez,N.,Melamed,S.,Paran,N.,Beth-Din,A.,Zvi,A. |
| **Singapore** | MT903342 | 2020-09-15 | 2019-05-XX | Singapore | IIb | Mauldin,M.R.,McCollum,A.M.,Nakazawa,Y.J.,Mandra,A.,Whitehouse,E.R.,Davidson,W.,Zhao,H.,Gao,J.,Li,Y.,Doty,J.,Yinka-Ogunleye,A.,Akinpelu,A.,Aruna,O.,Naidoo,D.,Lewandowski,K.,Afrough,B.,Graham,V.,Aarons,E.,Hewsor |
| **MT250197** | MT250197 | 2020-05-12 | 2019-XX-XX | Singapore | IIb | Yong,S.E.F.,Ng,O.T.,Ho,Z.J.M.,Mak,T.M.,Marimuthu,K.,Vasoo,S.,Yeo,T.W.,Ng,Y.K.,Cui,L.,Ferdous,Z.,Chia,P.Y.,Aw,B.J.W.,Manauis,C.M.,Low,C.K.K.,Chan,G.,Peh,X.,Lim,P.L.,Chow,L.P.A.,Chan,M.,Lee,V.J.M.,Lin,R.T.P.,Heng,M.K. |
| **UK_P3** | MT903345 | 2020-09-15 | 2018-09-XX | United Kingdom | IIb | Mauldin,M.R.,McCollum,A.M.,Nakazawa,Y.J.,Mandra,A.,Whitehouse,E.R.,Davidson,W.,Zhao,H.,Gao,J.,Li,Y.,Doty,J.,Yinka-Ogunleye,A.,Akinpelu,A.,Aruna,O.,Naidoo,D.,Lewandowski,K.,Afrough,B.,Graham,V.,Aarons,E.,Hewsor |
| **UK_P2** | MT903344 | 2020-09-15 | 2018-09-XX | United Kingdom | IIb | Mauldin,M.R.,McCollum,A.M.,Nakazawa,Y.J.,Mandra,A.,Whitehouse,E.R.,Davidson,W.,Zhao,H.,Gao,J.,Li,Y.,Doty,J.,Yinka-Ogunleye,A.,Akinpelu,A.,Aruna,O.,Naidoo,D.,Lewandowski,K.,Afrough,B.,Graham,V.,Aarons,E.,Hewsor |
| **USA_2021_MD** | ON676708 | 2022-06-03 | 2021-11-XX | USA | IIb | Gigante,C.M.,Myers,R.,Seabolt,M.H.,Wilkins,K.,McCollum,A.,Hutson,C.,Davidson,W.,Rao,A.,Blythe,D.,Li,Y. |
| **human/Taiwan/110-23** | ON918656 | 2022-07-05 | 2022-06-XX | Taiwan | IIb | Lin,J.-H.,Chiu,S.-C.,Huang,H.-I.,Huang,W.-L.,Fann,W.-B.,Hsieh,P.-Y.,Yang,J.-Y. |
| **human/Japan/Tokyo/T** | LC722946 | 2022-08-11 | 2022-07-XX | Japan | IIb | Kasuya,F.,Negishi,A.,Kumagai,R.,Hasegawa,M.,Fujiwara,T.,Miyake,H.,Nagashima,M.,Sadamasu,K. |
| **Germany/2023/ON/RK** | PP093715 | 2024-01-15 | 2023-11-XX | Germany | IIb | Brinkmann,A.,Kohl,C.,Schrick,L.,Michel,J.,Schaade,L.,Nitsche,A. |
| **Germany/2022/ON/RK** | OR463738 | 2023-08-27 | 2022-07-XX | Germany | IIb | Brinkmann,A.,Pape,K.,Kohl,C.,Schrick,L.,Michel,J.,Schaade,L.,Nitsche,A. |
| **60_D** | OR264443 | 2023-08-09 | 2022-08-22 | Australia | IIb | Taouk,M.L.,Steinig,E.,Taiaroa,G.,Savic,I. |
| **56_C** | OR264437 | 2023-08-09 | 2022-09-06 | Australia | IIb | Taouk,M.L.,Steinig,E.,Taiaroa,G.,Savic,I. |
| **Monkeypox/PT0158/2** | OP324475 | 2022-08-29 | 2022-07-07 | Portugal | IIb | Isidro,J.,Borges,V.,Pinto,M.,Sobral,D.,Santos,J.,Nunes,A.,Mixao,V.,Ferreira,R.,Santos,D.,Duarte,S.,Vieira,L.,Borrego,M.J.,Nuncio,S.,Lopes de Carvalho,I.,Pelerito,A.,Cordeiro,R.,Gomes,J.P. |
| **NY-NYCPHL-000389** | OQ468973 | 2023-02-26 | 2022-XX-XX | USA | IIb | Clabby,T.T.,Amin,H.S.,Wang,J.C.,Taki,F.,Su,M.,Rahat,A.,De La Cruz,N.,Olsen,A.,Thi,C.,Silver,S.,Akther,S.,Chowdhury,M.,Omoregie,E.,Hughes,S. |
| **ANT-LDSP-ANT-MPX-** | OQ261707 | 2023-01-22 | 2022-10-14 | Colombia | IIb | Betancur,I.I.B.,Velarde-Hoyos,C.-A.C.V.,Gomez,R.R.G.,Mercado-Reyes,M.M.R. |
| **ANT-LDSP-ANT-MPX-** | OQ957072 | 2023-05-17 | 2023-02-16 | Colombia | IIb | Betancur,I.I.B.,Velarde Hoyos,C.A.C.V.,Gomez,R.R.G.,Mercado-Reyes,M.M.R. |
| **ANT-LDSP-ANT-MPX-** | OP642395 | 2022-10-17 | 2022-09-07 | Colombia | IIb | Betancur,I.I.B.,Velarde-Hoyos,C.-A.C.V.,Gomez,R.R.G.,Mercado-Reyes,M.M.R. |
| **hMpxV/USA/IL-RIPHL** | OR574869 | 2023-09-26 | 2023-06-12 | USA | IIb | Green,S.,Kunstman,K.,Barbian,H.,Araujo Perez,F.,Kittner,A.,Bobrovska,S. |
| **Germany/2023/ON/RK** | PP182128 | 2024-01-28 | 2023-12-XX | Germany | IIb | Brinkmann,A.,Kohl,C.,Schrick,L.,Michel,J.,Schaade,L.,Nitsche,A. |
| **NY-NYCPHL-0001363** | PQ065598 | 2024-07-29 | 2024-07-XX | USA | IIb | Clabby,T.,Amin,H.S.,Taki,F.,Su,M.,Wang,J.C.,De La Cruz,N.,Olsen,A.,Thi,C.,Akther,S.,Chowdhury,M.,Omoregie,E.,Polanco,M.,Chen,X.,Siemetzki-Kapoor,U. |
| **Human/USA/CA-LACI** | PP918961 | 2024-06-23 | 2024-05-15 | USA | IIb | Heibeck,N.,Garrigues,J.,Green,N. |
| **USA_2022_CA006** | OP150924 | 2022-08-04 | 2022-06-XX | USA | IIb | Gigante,C.,Ventura,J.,Seabolt,M.H.,Zhao,H.,Wilkins,K.,McCollum,A.,Hutson,C.,Davidson,W.,Rao,A.,Nash,J.,Sheth,M.,Li,Y. |
| **human/Japan/Tokyo/I** | LC831698 | 2024-07-25 | 2024-03-03 | Japan | IIb | Higashi-Kuwata,N.,Okada,W.,Takahashi,K.,Nagashima,M.,Morioka,S.,Iwamoto,N.,Sadamasu,K.,Yoshimura,K.,Ohmagari,N.,Mitsuya,H. |
| **ROK-P53/04-2023** | OR855747 | 2023-12-04 | 2023-09-15 | South Korea | IIb | Chung,Y.-S.,Yi,H.,Choi,M.-M.,Kim,J.-W.,Lee,M.,Lee,S.E.,Sim,G.,Lee,J.H.,Shin,H.,Choi,C.-H. |
| **MPV/Human/USA/CA-** | OR643705 | 2023-10-11 | 2023-08-16 | USA | IIb | Garrigues,J.M.,Green,N.M. |
| **human/CHN/GDCDC_** | PP648206 | 2024-04-17 | 2023-06-10 | China | IIb | Li,B.,Zhao,W.,Shen,C. |
| **Germany/2024/ON/RK** | PP776546 | 2024-05-14 | 2024-02-XX | Germany | IIb | Brinkmann,A.,Kohl,C.,Schrick,L.,Michel,J.,Schaade,L.,Nitsche,A. |
| **PT0818/2024** | PP481196 | 2024-03-19 | 2024-01-30 | Portugal | IIb | Isidro,J.,Borges,V.,Pinto,M.,Sobral,D.,Santos,J.,Nunes,A.,Mixao,V.,Ferreira,R.,Santos,D.,Duarte,S.,Vieira,L.,Borrego,M.J.,Nuncio,S.,Lopes de Carvalho,I.,Pelerito,A.,Cordeiro,R.,Gomes,J.P. |
